# Parallel analysis of RNA ends reveals global microRNA-mediated target RNA cleavage in maize

**DOI:** 10.1101/2022.06.02.494554

**Authors:** Juan He, Chi Xu, Chenjiang You, Beixin Mo, Xuemei Chen, Lei Gao, Lin Liu

## Abstract

MicroRNAs (miRNAs) are endogenous 20-to 24-nucleotide (nt) noncoding RNAs that play important regulatory roles in many biological processes in eukaryotes. miRNAs modulate the expression of target genes at the post-transcriptional level by transcript cleavage or translational inhibition. Identification of miRNA target genes have been extensively investigated in *Arabidopsis* and rice, but an in-depth global analysis of miRNA-mediated target regulation is still lacking in maize. Here, we report a transcriptome-wide identification of miRNA targets by analyzing Parallel Analysis of RNA Ends (PARE) datasets derived from nine different tissues at five developmental stages of the maize (*Zea mays* L.) B73 cultivar. 246 targets corresponding to 60 miRNAs from 25 families were identified, including transcription factors and other genes. In addition, PARE analysis revealed that miRNAs guide specific target transcript cleavage in a tissue-preferential manner. Interestingly, primary transcripts of *MIR159c* and *MIR169e* were found to be cleaved by mature miR159 and miR169, respectively, indicating a positive-feedback regulatory mechanism in miRNA biogenesis. Moreover, several new miRNA-target gene pairs involved in seed germination were identified and experimentally validated. Our PARE analyses generated a wide and detailed miRNA-target interaction atlas, which provides a valuable resource for investigating the roles of miRNAs and their targets in maize.

## INTRODUCTION

Small RNAs (sRNAs) are 20-to 24-nucleotide (nt) noncoding RNAs of which two classes, microRNAs (miRNAs) and small interfering RNAs (siRNAs), have been identified in plants based on their precursor structures and mode of biogenesis (Chen and Rechavi, 2021; Yu et al., 2019; Axtell, 2013). miRNAs are endogenous small noncoding RNAs derived from *MIRNA* precursor transcripts with a characteristic stem-loop structure. After being precisely excised by the RNase III protein DICER-LIKE1 (DCL1) with the assistance of other RNA-binding cofactors, the miRNA duplexes are released and then 2′-O-methylated by the methyltransferase HUA ENHANCER 1 (HEN1) (Chen et al., 2002; Yu et al., 2005). Mature guide miRNA strands are subsequently incorporated into the ARGONAUTE1 (AGO1) effector to regulate the expression of their target genes by sequence complementarity at the post-transcriptional level (Ha and Kim, 2014; Rogers and Chen, 2013; Iwakawa and Tomari, 2015). In plants, miRNAs act as universal regulators in a wide range of developmental and physiological processes and responses to multiple stresses (Yu et al., 2019).

In plants, miRNAs have been reported to target a diverse range of regulatory genes, including a large number of transcription factors, suggesting that miRNAs have essential roles at the core of gene networks and participate in myriad regulatory pathways. For example, members of the conserved miR156 family take part in vegetative phase transition and flowering by modulating the expression of *SQUAMOSA PROMOTER BINDING PROTEIN-LIKE* (*SPL* or *SBP box*) genes (Wang, 2014). Furthermore, miR156 targets *Ideal Plant Architecture 1* (*IPA1*) /*SPL14*, which confers on rice an ideal plant architecture with reduced unproductive tiller numbers, increased lodging resistance and substantially enhanced grain yield, as well as improved disease resistance (Jiao et al., 2010; Liu et al., 2019). miR160 plays a critical role in plant growth and development by regulating target *AUXIN RESPONSE FACTOR* (*ARF*) genes. Repression of *ARF17* by miR160 has been shown to mediate hypocotyl growth in *Arabidopsis* (Mallory et al., 2005), while miR160 also regulates *ARF10* and *ARF16*, which are involved in seed germination (Liu et al., 2007; Liu et al., 2013). Moreover, in addition to targeting protein-coding genes, several miRNAs can target non-coding RNAs; for example, miR2118 triggers the cleavage of a long non-coding RNA (lncRNA) *Photoperiod-sensitive genic male sterility 1* (*PMS1T*), resulting in the generation of 21-nt phased small-interfering RNAs (phasiRNAs) involved in the regulation of male fertility under long-day conditions (Fan et al., 2016).

Maize (*Zea mays* L.) is one of the most productive staple crops worldwide, as well as a model plant for genetic study. In addition to being a critical source of food, feed, fuel and fiber, maize has tremendous genetic diversity, as revealed by high-throughput sequencing, which can be exploited to improve breeding and quality (Gore et al., 2009). Although miRNA functions are less well studied in maize than in *Arabidopsis* and rice, the vital roles of miRNAs in maize development are highlighted by the broad range of developmental defects in mutants of miRNA biogenesis, in specific miRNAs, or in miRNA target genes. The maize *fuzzy tassel* (*fzt*) mutant contains a mutation in *DCL1*, which results in reduced miRNA accumulation, and exhibits multifaceted developmental defects, including abnormal meristems, reduced plant height and altered leaf polarity (Thompson et al., 2014). The dominant mutant *Corngrass* (*Cg1*) caused by overexpression of miR156 retains a slender juvenile grass-like morphology and shows multiple further abnormalities in plant architecture (Chuck et al., 2007). miR156 targets members of the *SPL* family of transcription factors, including *teosinte glume architecture 1* (*tga1*) (Chuck et al., 2007) and *tasselsheath4* (*tsh4*) (Chuck et al., 2010). In addition, *tasselseed4* (*ts4*), which encodes miR172e, plays a key role in sex determination by repressing two *AP2*-like genes, *indeterminate spikelet1* (*ids1*) and *sister of indeterminate spikelet* (*sid1*) (Chuck et al., 2007; Chuck et al., 2008). miR164 inhibits the expression of the transcription factor gene *NAC1* and participates in the regulation of lateral root development (Li et al., 2012). In addition, miR166 contributes to the establishment of leaf polarity by regulating the Class III homeodomain/leucine zipper (*HD-ZIPIII*) family member *rolled leaf1* (*rld1*) (Juarez et al., 2004). Moreover, miR399 represses *PHOSPHATE2* (*PHO2*) and contributes to maize tolerance of low phosphate availability (Du et al., 2018).

Animal miRNAs mainly regulate target genes by translational repression (Karginov et al., 2010), while in plants miRNAs repress target mRNAs by both cleavage and translational repression (Addo-Quaye et al., 2008; German et al., 2008; Gregory et al., 2008). Many plant miRNAs recognize target mRNAs via nearly perfect sequence complementarity and guide the AGO1 protein to cleave the transcripts at the phosphodiester bond corresponding to miRNA nucleotides 10 and 11 (Chen, 2009). The identification of the targets of miRNAs is critical to the elucidation of their biological functions. Computational approaches have been established to predict miRNA targets on a genomic scale using base pairing rules; popular software packages include psRNAtarget (Dai et al., 2018; https://www.zhaolab.org/psRNATarget/) and Tarhunter (Ma et al., 2018; http://www.biosequencing.cn/TarHunter/). However, the predicted targets need to be experimentally validated due to high false positive rates. A powerful approach known as Parallel Analysis of RNA Ends (PARE) or degradome sequencing, which captures the monophosphated 5’-end of a cleaved mRNA 3’-fragment by ligation with an RNA adaptor (Addo-Quaye et al. 2008; German et al. 2008; Brian et al., 2008; Zhai et al., 2014), provides experimental evidence for miRNA-mediated target-RNA cleavage on a global scale. PARE has been successfully used to identify miRNA targets in *Arabidopsis* (Addo-Quaye et al. 2008; German et al. 2008), rice (Li et al., 2010; Zhou et al., 2010) and a few other plant species (Jeong, et al., 2013).

There are 371 miRNAs and miRNA*s forming 73 miRNA families in maize based on data from miRbase (Kozomara et al., 2019) and miRNEST (Szczesniak et al., 2014). Compared to the well-characterized miRNA targets in *Arabidopsis* and rice (Addo-Quaye et al. 2008; German et al. 2008; Tang and Chu, 2017), only a few miRNA targets have been identified in maize (Chuck et al., 2007; Chuck et al., 2010; Chuck et al., 2008; Li et al., 2012; Juarez et al., 2004; Du et al., 2018). To comprehensively identify transcriptome-wide miRNA targets in maize, we constructed PARE libraries using nine maize tissues at five developmental stages in this study. Different subsets of miRNA target genes were identified in different tissues with the highest number of targets in tassel and the lowest in silk at the reproductive stage, which roughly corresponds to the numbers of miRNAs expressed in these tissues. In addition, a series of novel targets for conserved miRNAs and monocot-specific miRNAs were identified. Moreover, this study revealed that some miRNAs with similar but distinct sequences, such as miR156/miR529 and miR159/miR319, target different subsets of genes in different tissues, implying their functional divergence in maize. Finally, differentially expressed miRNAs and their target genes involved in seed germination were identified and experimentally validated. This study provides a comprehensive atlas of miRNAs and their target genes in various tissues and from different developmental stages, which should enable further studies of various biological processes regulated by miRNAs in maize.

## RESULTS

### Preparation of PARE libraries from different tissues of maize

Nine different tissues, including dry seed (seeds at the dent stage), germinating seed, shoot and root at the V1 stage (collar of first leaf visible at the vegetative stage), stalk at the V3 stage (collar of third leaf visible at the vegetative stage), as well as leaf, tassel, silk and ear at the R1 stage (silk just visible outside the husks at the reproductive stage), of the maize (*Zea mays*) inbred line B73 were collected for RNA extraction and construction of PARE libraries. For each sample, a total of over 30 million raw sequencing reads with adaptors was obtained, among which ∼20 million 20-nt reads representing ∼70% of the total clean reads could be mapped to annotated transcripts of maize B73 (Supplemental Dataset 1). At least two replicates with high correlation coefficients were obtained for each tissue (Supplemental Figure S1A). We have published RNA-seq and sRNA-seq data for some of the tissues previously (He et al., 2019) and these were used in the analysis together with the PARE datasets. sRNA-seq and RNA-seq libraries for other tissue types were constructed and sequenced in this study (Supplemental Figure S1B and C, Supplemental Dataset 2 and 3). About 16 million uniquely mapped reads per sample for RNA-seq and more than 4 million mapped reads for sRNA-seq were obtained for each sample (Supplemental Dataset 2 and 3). At least two biological replicates were conducted for each sample with high correlation coefficients.

### PARE analysis provides a global view of miRNA-directed target cleavage in maize

Transcripts sliced by sRNAs should have distinct peaks of PARE reads at the predicted cleavage sites – the 10^th^ nt counting from the 5’-end of each miRNA (Addo-Quaye et al., 2008; German et al., 2008). The CleaveLand pipeline (Addo-Quaye et al., 2009; Brousse et al., 2014) was used to identify 5’-ends generated by miRNA-guided cleavage in the PARE datasets. The identified transcripts were grouped into five categories (categories 0-4) based on read abundance at the target site relative to reads along the entire transcript, as reported previously (Brousse et al., 2014). Transcripts found in category 0 had the highest credibility as real miRNA targets, because PARE reads are more abundant at the predicted target site than the rest of the transcript; about half the targets in each maize tissue fell into this category (Figure 1A). Targets for which PARE signatures indicative of miRNA-mediated cleavage were among the most abundant reads over the length of the transcript belonged to category 1; no more than 26 targets were found in this category (Figure 1A; Supplementary Dataset 4). As to category 2, the abundance of PARE reads indicative of miRNA-mediated cleavage was above the median, but below the maximum on the transcript. As shown in Figure 1A, 16 to 45 targets from different tissues were in this category. Category 3 encompassed transcripts with more than one read at the 5’-end of a slicing remnant, but the abundance was below or equal to the median of reads across the entire transcript. Category 4 covered those transcripts with only one read at the miRNA-mediated cleavage site. In the nine maize tissues tested, 6-18 targets fell into category 3 and 3-15 fell into category 4 (Figure 1A; Supplemental Dataset 4). Genes in categories 3 and 4 might not be true miRNA target genes.

**Figure 1.**
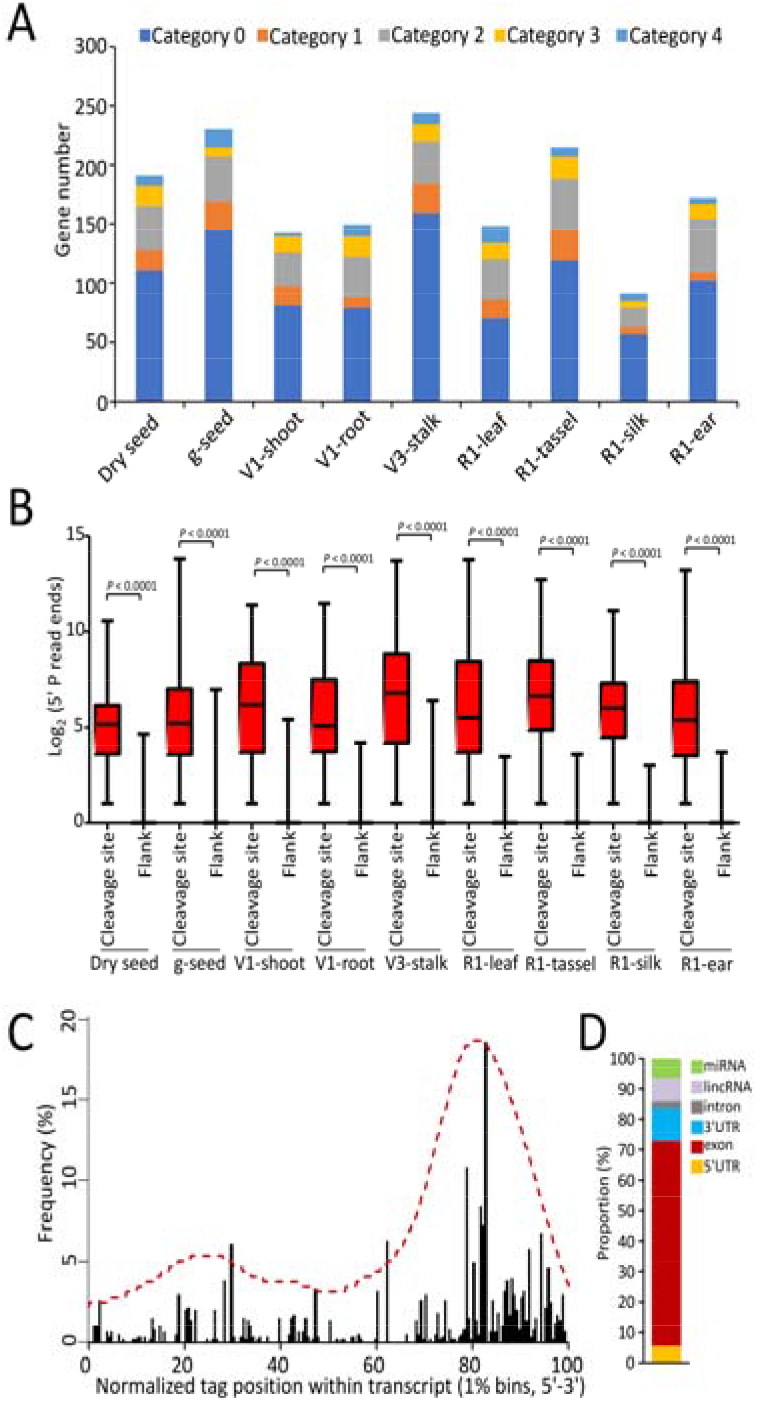
Identification of target genes of maize miRNAs by PARE sequencing. A, Gene numbers of target genes belonging to category 0-4 in different maize tissues. g-seed: germinating seed, V1: vegetative stage 1, V3: vegetative stage 3, R1: reproductive stage 1. B, Accumulation of 5’-P read ends at miRNA target sites. Flank: the 100-nucleotide flanking region of the cleavage site. C, Histogram displaying the 5’-P position of degradome tags of miRNA target genes relative to normalized transcript position. Tags were counted in 1% bins. D, Proportion of miRNA cleavage sites in the genome.

The miRNA cleavage sites in category-0 targets were represented by read counts in our PARE data up to several thousand-fold greater than those in the 100 nucleotides flanking these regions (local cleavage efficiency) and several hundred-fold more than the average coverage of 5’-P reads along the entire transcript (global cleavage efficiency) (Figure 1B; Supplemental Dataset 5). The miRNA-directed cleavage efficiency was much higher in V3-stalk, R1-tassel and V1-shoot than in other tissues (Figure 1B), consistent with the fact that these three tissues had the highest numbers of abundant miRNAs (Supplemental Figure S2). The PARE tags for miRNA targets were biased toward the 3’-ends of annotated transcripts (Figure 1C) and exon regions of genes had the most target sites, followed by the 3’-UTR, lncRNA and 5’-UTR regions (Figure 1D).

### PARE identifies differential cleavage by miRNAs in different maize tissues

Our sRNA-seq data identified a total of 60 miRNAs from 25 families that were represented in at least one of the nine maize tissues spanning five developmental stages (Supplemental Figure S2A). The expressed miRNAs were most enriched in R1-tassel, V1-shoot and V3-stalk, implying active regulation by miRNAs in these tissues. Moreover, dozens of target genes for these expressed miRNAs were identified from different maize tissues (Supplemental Figure S3A). Relatively high numbers of target genes were found in germinating seed, R1-tassel, V3-stalk and V1-shoot, while fewer targets were found in dry seed and R1-silk (Supplemental Figure S3A). These results indicated that miRNA-directed cleavage was probably more active in tissues undergoing active developmental processes. Target genes for miRNAs were identified in different maize tissues, especially for those in category 0, which represent the most likely candidates (Supplemental Figure S3B). For example, more miR156 targets were found in V1-shoot, V1-root and V3-stalk, which are both young and vegetative tissues (Supplemental Figure S3B), implying a function of miR156 in juvenile maize growth, similar to its role in other plant species like *Arabidopsis* (Wu et al., 2009). Targets in category 0 for miR529 were mainly identified in R1-tassel (Supplemental Figure S3B), which was consistent with the specific and high level of expression of miR529 in R1-tassel (Supplemental Figure S2A). It is worth noting that a high level of miR529 was also observed in germinating seed (Supplemental Figure S2A). Accordingly, three targets of miR529 in category 0 were identified in germinating seed (Supplemental Figure S3B, Supplemental Dataset 4), suggesting that miR529 may be involved in the regulation of seed germination. Similarly, target genes of miR319 were primarily found in germinating seed (Supplemental Figure S3B), suggesting a specific role of miR319 in seed germination. Five common target genes, corresponding to miR166, miR159, miR171, miR528 and miR444, were found in all tissues examined, implying that they function in a multitude of developmental and physiological processes in maize (Figure 2A and 2B). Numbers of tissue-specific targets was also identified from our PARE analysis, including a diverse range of regulatory genes, such as transcription factors, as well as R (resistance) genes and genes encoding metabolic enzymes (Figure 2A). The distinct patterns of miRNA-directed cleavage in different maize tissues signify the complex regulatory mechanisms orchestrated by miRNAs during development.

**Figure 2.**
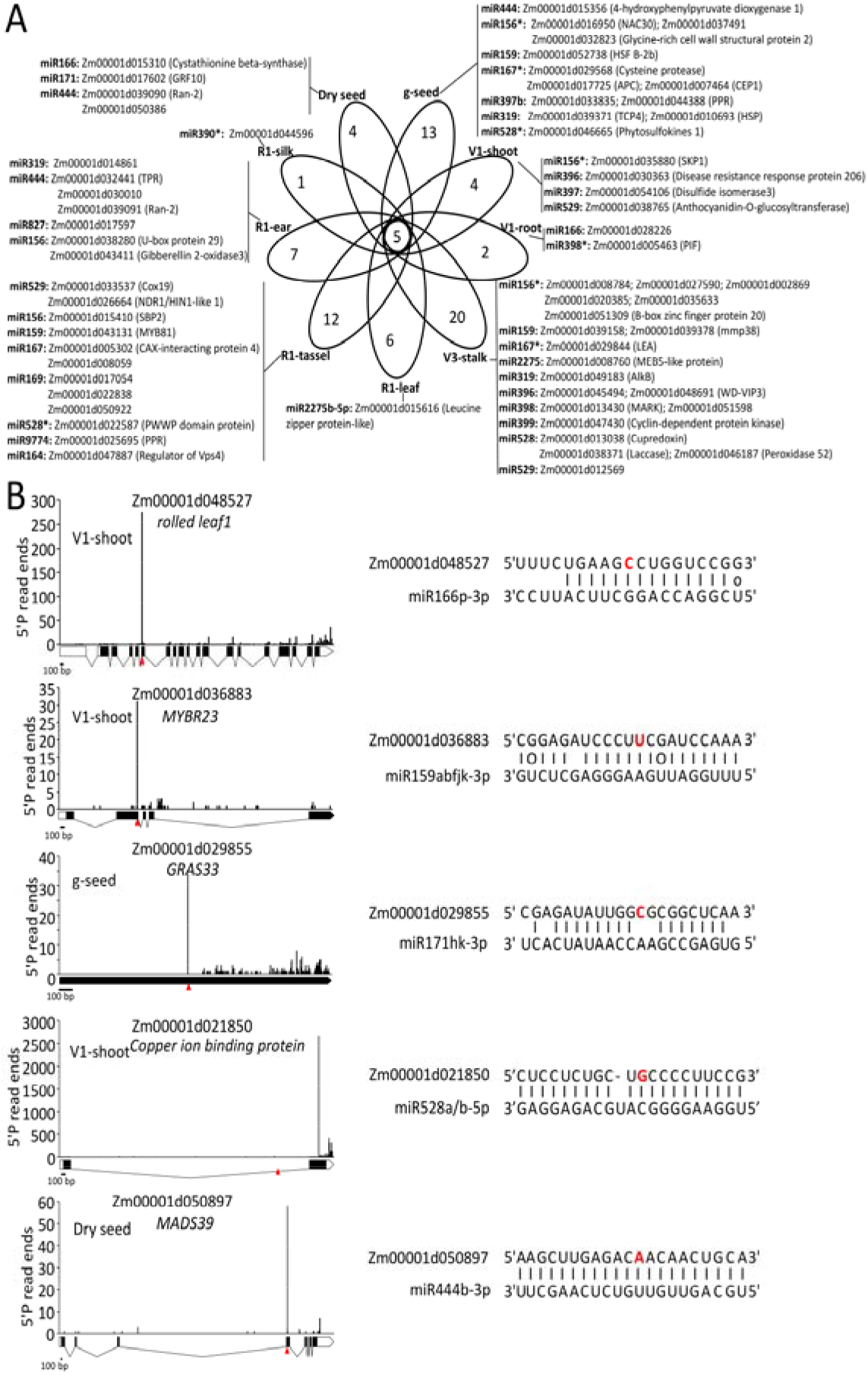
Summary of miRNA target genes identified by PARE-seq in different maize tissues. A, Venn diagrams for identified target genes of maize miRNAs by PARE-seq. V1: vegetative stage 1, V3: vegetative stage 3, R1: reproductive stage 1. B, T-plots and alignment of the common target genes of miRNAs identified in all maize tissues of analyzed. The red nucleotides indicate the 5’-P end for the residues miRNA target genes detected in the PARE analysis and the respective arrowheads show the cleavage sites. In the gene model, the white boxes illustrate the untranslated regions (UTRs), the black boxes show the coding regions(CDS) and the black lines indicate the introns of the genes.

### Tissue-specific targeting of miR159/miR319 and miR156/miR529 in maize

Two pairs of miRNAs, miR156/miR529 and miR159/miR319, show a high degree of sequence similarity between the members of each pair, which is conserved across plant species (Morea et al., 2016; Palatnik et al., 2007; Xie et al., 2021). In maize, there were 16-17 identical nucleotides between the two miRNAs of each pair (Figure 3A and Figure 4A), making their target genes hard to distinguish by computational prediction.

**Figure 3.**
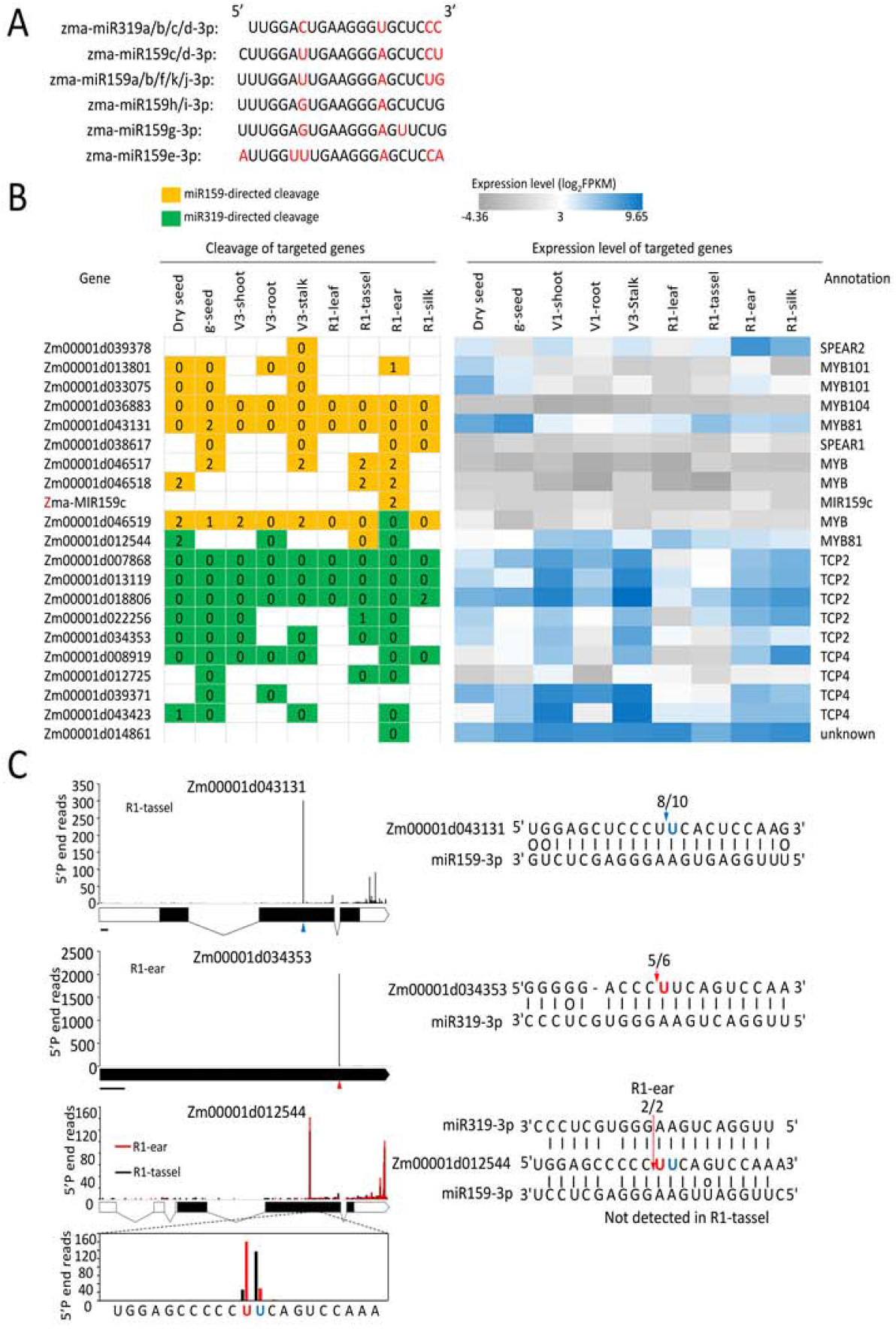
Tissue-specific targeting of miR159 and miR319 in different tissues of maize. A, Sequences alignment of maize miR159 and miR319. The different nucleotides between sequences are shown in red. B, Target genes of miR159 and miR319 across nine different tissues in maize. The diagram on the left shows the targeting patterns of miR159 and miR319 in different tissues. Numbers in the colored boxes indicate to which categories the relevant target genes belong. The diagram on the right illustrates the expression patterns of those target genes, g-seed: germinating seed. V1: vegetative stage 1, V3: vegetative stage 3, R1: reproductive stage 1. C, Target plots (T-plot) and 5’-RACE verification of representative targets of miR159 and miR319 in maize. The red and blue nucleotides indicate the 5’-P end for the residues of miR319 and miR159 target genes detected in the PARE analysis, respectively. The respective arrowheads or arrows show the cleavage sites. The numbers above the alignments indicate the data from 5’-RACE confirmation. In the gene model, the white boxes illustrate the UTRs, the black boxes show the CDS and the black lines indicate the introns of the genes.

**Figure 4.**
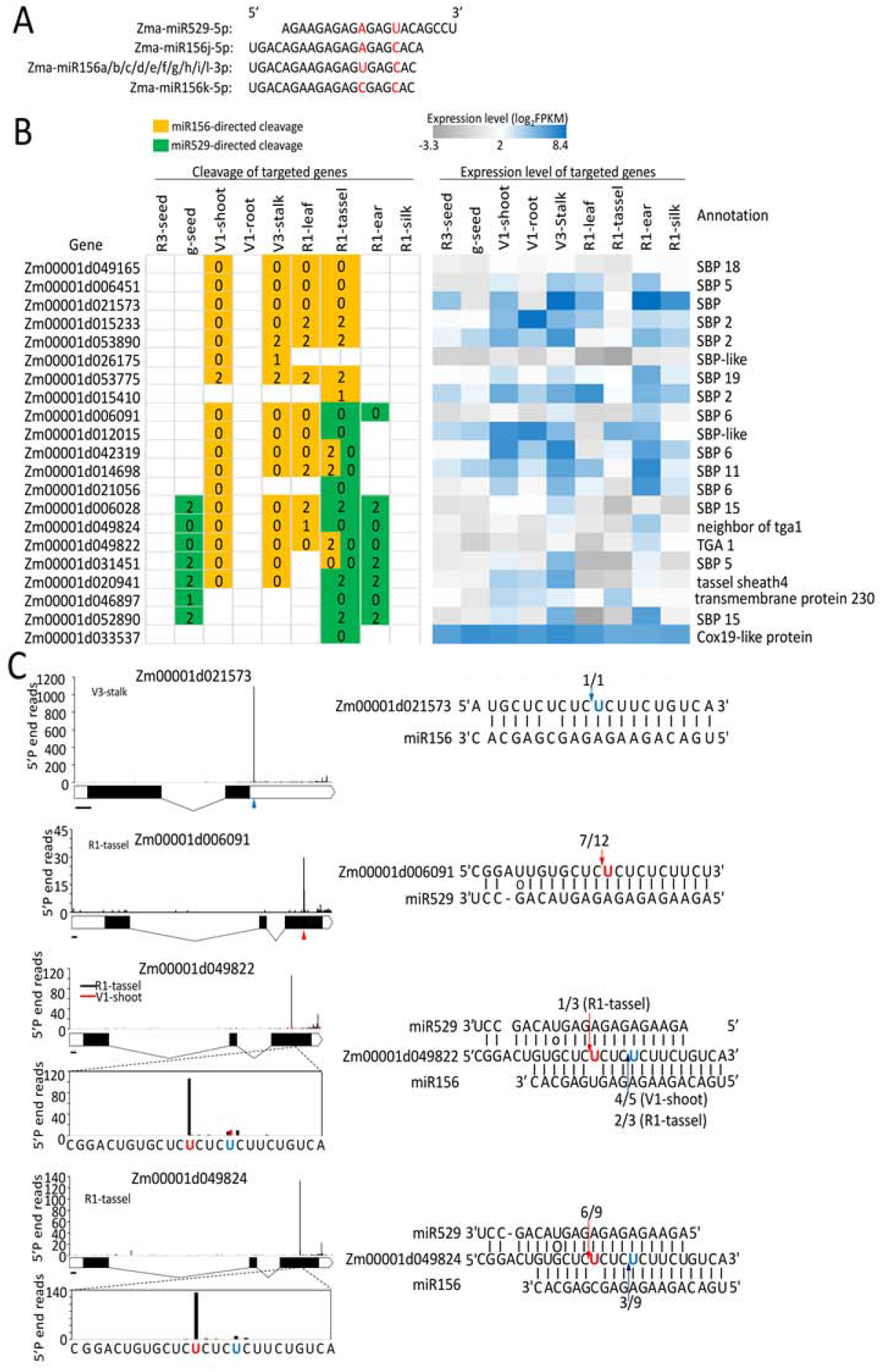
miR156 and miR529 have distinct subsets of targets in different maize tissues. A, Sequence alignment of maize miR156 and miR529. The different nucleotides between sequences are shown in red. B, Target genes of miR156 and miR529 in different maize tissues. The diagram on the left shows the targeting patterns of miR156 and miR529 in different tissues. Numbers in the colored boxes indicate to which categories the relevant target genes belong. The diagram on the right illustrates the expression patterns for those target genes, g-seed: germinating seed. V1: vegetative stage 1, V3: vegetative stage 3, R1: reproductive stage 1. C, Target plots (T-plot) and 5’-RACE verification of representative targets of miR156 and miR529 in maize. The red and blue nucleotides indicate the 5’-P ends for the residues of miR529 or miR156 target genes detected in the PARE analysis, respectively. The respective arrowheads or arrows show the cleavage sites. In the gene model, the white boxes illustrate the UTRs, the black boxes show the CDS and the black lines indicate the introns of the genes.

*MIR159* and *MIR319* genes, which evolved from a common ancestor, are highly conserved across land plants (Li et al., 2011). Moreover, *MIR319* gene seems to emerge earlier than *MIR159* in the genome of angiosperm, which accords with the more specialized target spectrum of miR159 than miR319 (Palatnik et al., 2007). In *Arabidopsis*, miR159 targets *MYB* mRNAs, while miR319 predominantly regulates *TCP* mRNAs, but can also act on some *MYB* mRNAs. The targets of miR159 and miR319 in maize were generally similar to those in *Arabidopsis* (Figure 3B). Some novel target genes were identified in our PARE datasets. Besides *MYB* genes, two new target genes named *SPOROCYTELESS-like EAR-containing 1* (*SPEAR1*) and *SPEAR2*, which are reported to act as transcriptional repressors in *Arabidopsis* and rice (Chen et al., 2014; Ren et al., 2018), were identified as targets for miR159 only. *SPEAR1* was found to be cleaved in maize female tissues including R1-ear and R1-silk as well as germinating seed and V3-stalk, while *SPEAR2* was only found in V3-stalk (Figure 3B). Interestingly, the cleavage of *MYB81* (*Zm00001d012544*) by miR159 only occurred in the male tissue tassel, while two *MYB101* (*Zm00001d013801* and *Zm00001d033075*) cleavages mainly occurred in vegetative tissues; cleavage of *MYB104* (*Zm00001d036883*) and *MYB81* (*Zm00001d043131*) was identified in all developmental stages examined in maize (Figure 3B and 3C). On the other hand, miR319 targets were mainly confined to the *TCP* genes, just like in *Arabidopsis*. Two *MYB* (*Zm00001d046519* and *Zm00001d012544*) mRNAs were targeted by both miR159 and miR319 (Figure 3B) in different tissues, implying functional specialization for these two miRNAs in maize development. Targets of miR159 and miR319 were randomly selected for validation by 5’-RACE, and the results further confirmed the PARE analysis (Figure 3C).

miR156 and miR529 are another pair of sequence-similar miRNAs. miR156 is highly conserved in almost all embryophytes while miR529 seems only to exist in monocots and some bryophytes (Cuperus et al., 2011; Xie et al., 2021). miR156 is a well-studied miRNA and plays important roles in the juvenile phase of many plant species (Wu et al., 2009; Chuck et al., 2007). A previous study showed that one target of maize miR156 was *teosinte glume architecture* (*tga1*), a gene known to have a role in the domestication of maize from teosinte (Chuck et al., 2007). In our study, we found interesting cleavage patterns of *TGA1* mRNA in different maize tissues by miR156 or miR529. In V1-shoot, V3-stalk and R1-leaf, *TGA1* mRNA was only targeted by miR156, while in germinating seed and R1-ear, it was only cleaved by miR529 (Figure 4B and 4C). Furthermore, *TGA1* mRNA can be targeted by both miR156 and miR529 in R1-tassel, but the cleavage products of miR529 were predominant. These results indicated that the expression of *TGA1* was coordinately regulated by miR156 or miR529 in different tissues to maintain its low expression level in most tissues and thereby to avoid the grass-like phenotype. Like *TGA1*, other target mRNAs of miR156 or miR529 also showed the tissue-specific cleavage patterns. Among them, mRNAs specifically targeted by miR156 mostly belonged to the *SQUAMOSA promoter-binding protein* (*SBP)* family, while two out of three miR529-specific targets were mRNAs of a transmembrane protein and a Cox19-like protein (Figure 4B). These results imply the existence of a sophisticated regulatory network involving miR156 and miR529 in maize developmental processes. Targets of miR156 and miR529 were randomly selected for validation by 5’-RACE, and the results confirmed the PARE analysis (Figure 4C).

### Self-cleavage of certain *MIRNA* primary transcripts in specific maize tissues

Some PARE tag signatures corresponding to *MIRNA* primary transcripts (pri-miRNAs) were found in our libraries. Pri-miRNAs with a major signature located where the mature miRNAs began were commonly observed (Supplemental Figure S4-6); these signatures likely represent intermediate products in pri-miRNA processing by DCL1 (Yu et al., 2018). It is interesting that whether mature miRNAs were located in the 5’ or 3’ arms of their respective pri-miRNAs, the PARE tag signatures were mostly found at the start of the mature miRNAs (Supplemental Figure S4-6). Some like pri-miR398b and pri-miR399e had high t-signatures corresponding to the center of their mature miRNAs or miRNA stars (Supplemental Figure S4), indicating that the pri-miRNAs may be targets of their own mature miRNAs. Although most such t-signatures did not correspond precisely to the position of the 10^th^ nucleotide of the mature miRNAs, except two cases. Pri-miR159c had abundant signatures at the expected site of cleavage by miR159a/b/f/j/k, specifically in R1-ear, with no other abundant signatures detected in the vicinity (±30 nt) (Figure 5A). Another one is pri-miR169e, showed high signatures exactly matching the 10^th^ nt from the 5’-end of miR169m/n/q (Figure 5B), and the cleavage was only being found in germinating seed and R1-ear of maize (Figure 5B). Moreover, we used 5’-RACE experiments to confirm the self-cleavage of pri-miR159c and pri-miR169e (Figure 5C). Previous study showed that pri-miR172b can be cleaved by its own mature miRNA in *Arabidopsis* (German et al., 2008), but no corresponding PARE signatures were found in the pri-miR172 members detected in our study, suggesting that self-cleavage of pri-miR172 probably does not occur in maize (Supplemental Figure S6). These results indicate that miRNA pathways can have complex feedback regulatory mechanisms that are specific to particular plant species and tissues.

**Figure 5.**
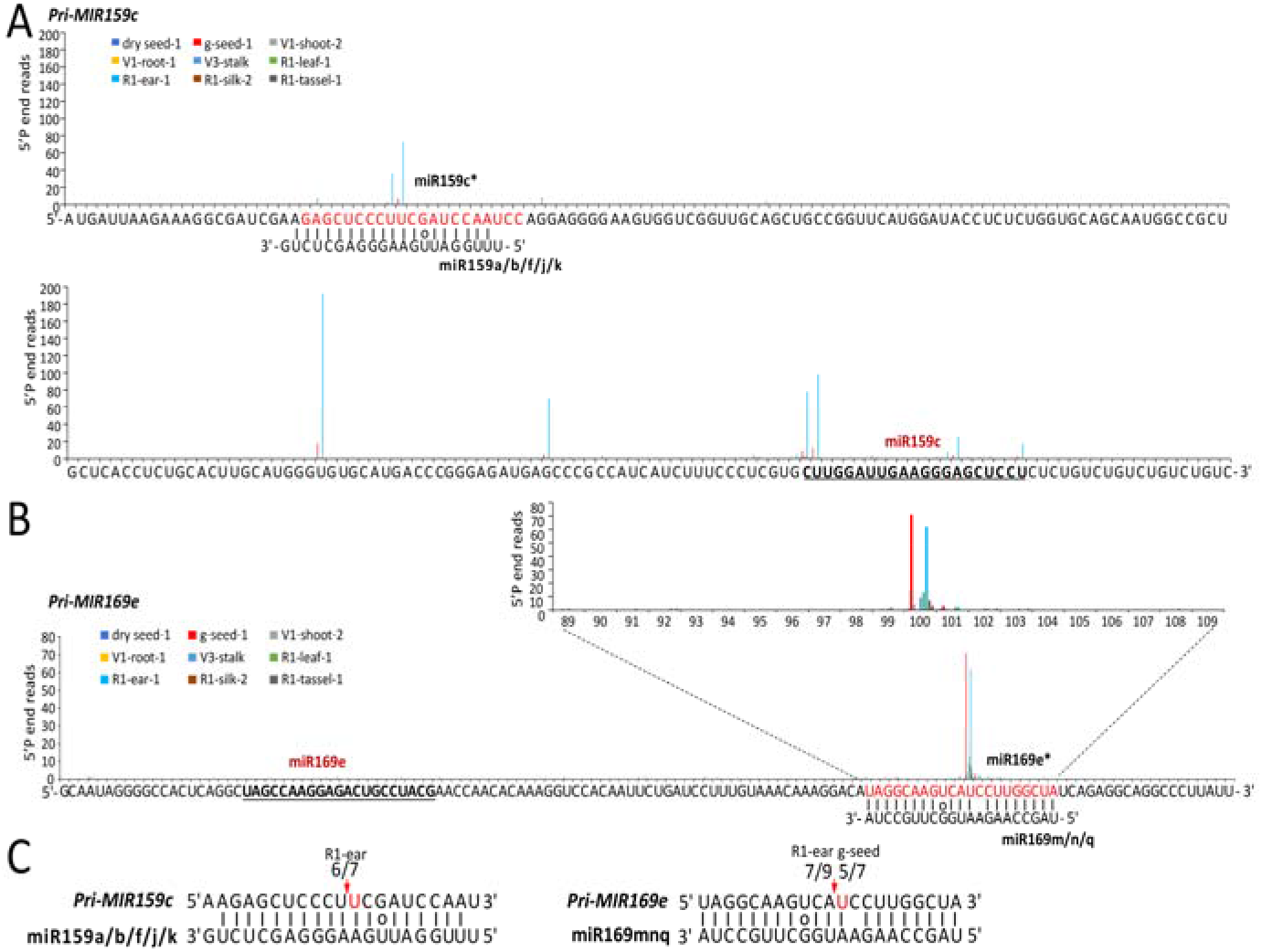
miRNAs primary transcripts can be targeted by their own mature miRNAs. A, pri-MIR159c has dominant signatures corresponding to mature miR159a/b/f/j/k at position 10. The mature miRNAs are underlined and miRNA*s were in red. g-seed: germinating seed. V1: vegetative stage 1, V3: vegetative stage 3, R1: reproductive stage 1. B, pri-MIR169e has dominant signatures corresponding to mature miR169m/n/q at position 10. C, Validation of miR159/miR169-guided cleavage of pri-MIR159c/pri-MIR169e by conventional S’-RACE. The red arrows indicate the cleavage sites and the nucleotides indicate the 5’-P end for the residue of target genes. The numbers above the alignments indicate the data from S’-RACEconfirmation.

### miRNAs and target genes are involved in the regulation of maize seed germination

Dormant seeds can germinate under appropriate temperature and humidity conditions. Recent reports have implicated miRNAs in the regulation of seed germination (Weitbrecht et al., 2011). Previous study confirmed the presence of 115 known miRNAs and 167 novel miRNAs at the early stage of seed germination in maize (Wang et al., 2011). However, differentially expressed miRNAs during maize seed germination have not been identified, and the target genes regulated by these miRNAs are still largely unknown.

To identify the key miRNA-target pairs involved in maize seed germination, we constructed sRNA-seq and corresponding PARE-seq libraries using dry seed and germinating seed. We compared the abundance of miRNAs in dry and germinating seed to identify differentially expressed miRNAs. Eight miRNAs belonging to five different miRNA families (miR159, miR166, miR396, miR528 and miR529) were identified (Figure 6A), which were quite different from the differentially expressed miRNAs identified in germinating seeds of *Arabidopsis* (Sarkar et al., 2018). All the miRNAs identified were expressed at a higher level in germinating seeds than in dry seeds, suggesting they have important roles in the induction of seed germination. For the miRNAs upregulated during germination, we identified target mRNAs that satisfied the following three criteria: 1) the targets contained sequences complementary to the corresponding miRNAs, 2) the cleavage signature of the mRNAs was higher in germinating seeds than in dry seeds, and 3) each target showed an approximately anti-correlated expression profile to that of the corresponding miRNA. Three differentially expressed miRNAs were selected for validation by northern blotting (Figure 6B). For miR159, we identified a new target gene *SPEAR1* (Figure 6C), which is a key repressor of sporogenesis in flowering plants (Chen et al., 2014; Ren et al., 2018). Two target genes for miR166 were identified, among which *rolled leaf 1* (*rld1, Zm00001d048527*) was previously reported to be a target gene of this miRNA (Juarez et al., 2004). The other unknown new target gene *Zm00001d026307* encodes a DAG (diacylglycerol) protein that may function in chloroplast development (Chatterjee et al., 1996; Luo et al., 2017) (Figure 6D). *Neighbor of tga 1* (*not1*) was found to be specifically cleaved during seed germination by miR529 (Figure 6E), implying a potential regulatory role for miR529 in germination.

**Figure 6.**
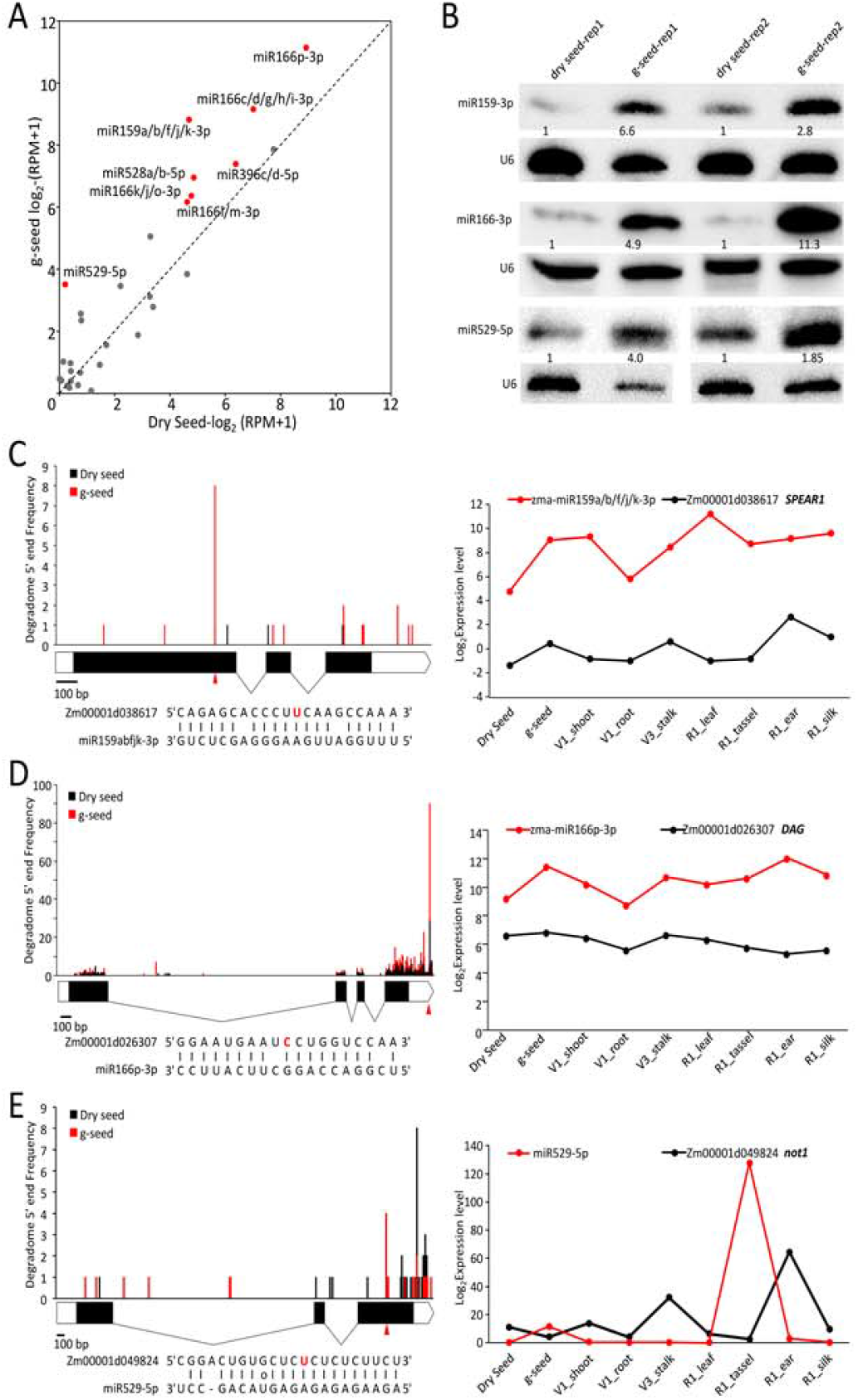
Novel targets of miRNAs involved in seed germination. A, miRNAs differentially expressed during seed germination. Red dots indicate miRNAs upregulated during germination. B, Validation of differential expression of miRNAs by northern blotting. Two biological replicates were conducted for each blot. U6 served as a loading control. Numbers below the blots indicate the abundance of the miRNAs relative to the dry seed control. C-E, The target plots (T-plots) of miR159 (C), miR166 (D) and miR529 (E) target genes show the abundances of PARE tags along the full length of the target mRNA sequences. The alignments show the miRNA with a portion of its target sequence. The red and blue nucleotides indicate the 5’-P ends for the residues of target genes detected in the PARE analysis. The arrowheads or arrows show the cleavage sites. The line charts show the expression patterns of miRNA and target genes in different maize tissues. The Y-axis indicates RPM (reads per million mapped reads) for miRNAs and RPKM (reads per kilobase per million mapped reads) for genes, g-seed: germinating seed. V1: vegetative stage 1, V3: vegetative stage 3, R1: reproductive stage 1.

To confirm that the three miRNAs direct the cleavage of their respective target transcripts, we performed a dual-luciferase transient expression assay in tobacco leaves. The sequences of pre-miR159, pre-miR166 and pre-miR529, driven by the cauliflower mosaic virus 35S promoter, were used as effectors, and the corresponding target sites or mutated target sites fused with luciferase (*LUC*) gene were used as reporters (Figure 7A). The results showed that the presence of the miRNAs significantly repressed *LUC* activity, which represents the expression levels of target genes, but the suppression was abolished when the target sites were mutated (Figure 7B and C). These results further confirmed the interactions between the three miRNAs and their corresponding targets during maize seed germination.

**Figure 7.**
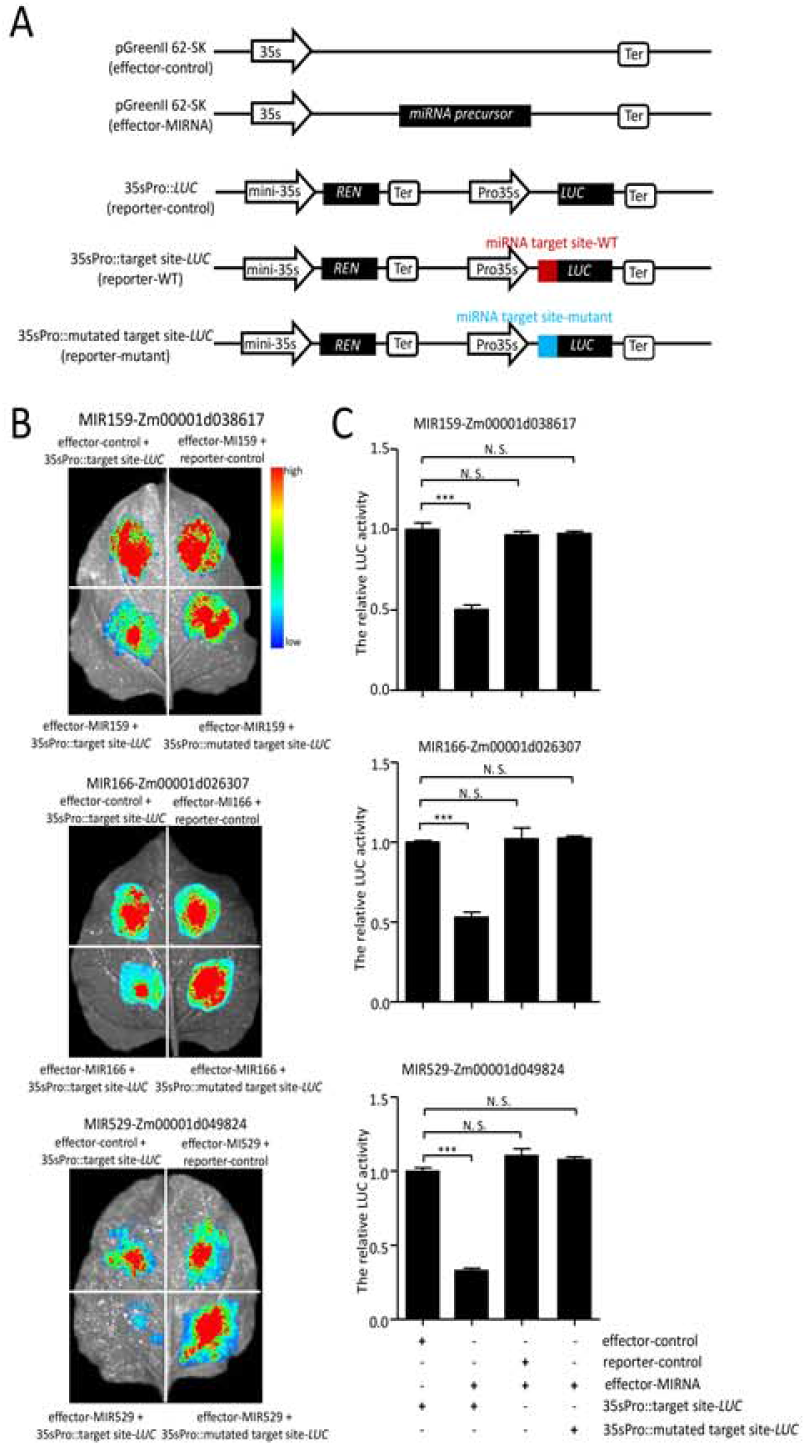
Dual-luciferase transient expression assay in tobacco leaves. A, Schematic diagrams indicating the constructs used in the dual-luciferase transient expression assays. B, Tobacco leaves co-transformed with different combinations of miRNA and target genes. C, Luciferase activity in the transient expression assays shown in B. Values are presented as means ± SD (n = 3). ***,p< 0.001; N.S., not significant.

## DISCUSSION

miRNAs participate in various developmental and physiological processes by targeting a diverse range of regulatory genes in plants (Chen and Rechavi, 2021). Large numbers of miRNAs were identified in maize by sRNA sequencing (Zhang et al, 2009), and many of them were revealed to be differentially expressed in various tissues and developmental stages (Liu et al., 2014; Xing et al., 2017), indicating their diverse functions in fine-tuning maize growth and development. In this study, we identified a comprehensive miRNA-target regulatory network from nine maize tissues spanning five different developmental stages. A total of 246 target genes were identified for 60 miRNAs from 25 families. Among them, several novel tissue-specific targets were revealed, for example *SPEAR2* targeted by miR159 in V3-stalk (Figure 3B), and *Cox19-like* gene targeted by miR529 in R1-tassel (Figure 4B). Furthermore, two miRNA primary transcripts, pri-miR159c and pri-miR169e, were found to be cleaved by mature miRNAs from their own family (Figure 5). In addition, several miRNA-target pairs essential to maize seed germination were identified and experimentally verified. These results indicate that PARE sequencing is an efficient strategy for identifying miRNA targets in maize.

As observed in *Arabidopsis* and rice, more than half of miRNA targets identified in maize encode transcription factors or have a role in transcriptional regulation (Jones-Rhoades et al., 2006; Zhou et al., 2010; Supplemental Figure S7). Furthermore, GO analysis of biological function for these target genes showed that they are mainly involved in metabolic and biosynthetic processes (Supplemental Figure S7). In addition, many other types of target genes were also identified. For example, three cupredoxin and laccase genes were identified as targets of miR528, while a cyclin-dependent protein kinase gene was revealed to be a candidate target of miR399. These results indicate that miRNAs and their targets function at the core of gene networks and play critical roles in a diverse range of pathways in plants.

Interestingly, some miRNA-target pairs were found to be regulated through auto-regulatory loops. For instance, miR168 targets AGO1-mediated cleavage of *AGO1* mRNA, and this feedback loop helps to maintain AGO1 homeostasis, which is critical for the proper functioning of the miRNA pathway. The miR168-*AGO1* module has been demonstrated to play an intricate role in regulating antiviral immunity, flowering time and tiller development by modulating multiple miRNAs (Varallyay et al., 2010; Wang et al., 2021 and Vaucheret et al., 2004). In addition, pri-miR172b was found to be cleaved by mature miR172 in *Arabidopsis* and was regulated by a feedback loop (German et al., 2008). Similarly, in this study we found that miR159 and miR169 could also target their own primary transcripts and cause self-cleavage (Figure 5). These findings indicate the presence of complicated regulatory mechanisms in miRNA biosynthesis and turnover. It is possible that there is competition between the folding process and the self-cleavage of miRNA precursors, which would define a homeostatic regulatory loop controlling miRNA biogenesis.

Phylogenetic studies have indicated that the evolution of *MIR* genes is in constant flux (Chavez et al., 2014). Some new paralogous *MIR* genes were generated by duplications of existing genes and then underwent functional diversification during evolution. Two clear cases are the miRNA pairs, miR159/miRNA319 and miR156/miR529, which show a high degree of sequence similarity within each pair across many plant species. *MIR159* and *MIR319* originated in the ancestor of seed plants and diverged into two different clades (Li et al., 2011). The *MIR319* clade diverged earlier than the *MIR159* clade in angiosperms, which is in accordance with the finding that miR319 has a more extensive range of targets than miR159. In *Arabidopsis* miR159 only regulates *MYB* mRNAs, while miR319 can target both *TCP* and *MYB* mRNAs (Palatnik et al., 2007). In our study, in addition to *MYB* genes, two new miR159 targets, namely *SPEAR1* and *SPEAR2*, were identified (Figure 3B). *SPEAR* genes act as transcription repressors and play crucial roles in the development of germ-line cells (Chen et al., 2014; Ren et al., 2018; Li et al., 2019). Dissecting the potential functions of *SPEAR1* and *SPEAR2* in maize will expand our understanding of the regulatory role of the miR159 family. The sequences of miR156 and miR529 are also very similar to each other, but they evolved independently and exhibit distinct evolutionary patterns (Xie et al., 2021). miR156 is conserved in all terrestrial plant species and is highly expressed during the vegetative stage, while miR529 is almost absent in core eudicots and is preferentially expressed at the reproductive stage (Morea et al., 2016; Xie et al., 2021). Consistent with previous studies, we found that maize miR529 was abundant in the tassel and at relatively low levels in the germinating seed and R1-ear, and was not detectable in other tissues (Supplemental Figure S2A). *SPL* genes are the major targets of miR156 and miR529 in many plant species (Morea et al., 2016). Interestingly, in addition to *SPL* targets, two novel targets specifically cleaved by miR529 were revealed by our PARE analysis, i.e., a *Cox19-like* gene from tassel and a gene of unknown function encoding a transmembrane protein in tassel and ear (Figure 4B). Further investigation of the functions of these two novel targets would help to elucidate the role of miR529 in the regulation of the development of maize reproductive tissues.

In summary, we have provided a comprehensive genome-wide characterization of miRNA targets by analyzing high-throughput PARE datasets of various maize tissues. This study will lead a clear understanding for the role of miRNAs and their targets during maize development. Although PARE sequencing is a powerful technique, no credible targets were found for some miRNAs in our study. A possible explanation of this observation is that some miRNAs are expressed in specific tissues or under different biotic or abiotic stresses, and thus were not detected in our study. The second possibility is that the targets were repressed at the translational level, which would not be detected by PARE sequencing. It is also possible that some targets were regulated by unknown miRNAs or siRNAs.

## MATERIALS AND METHODS

### Plant materials and growth conditions

All plant tissues were collected from maize (*Zea mays* ssp. *mays*) B73 plants, which were cultivated in a phytotron at 25°C and 65% relative humidity under a 14 h/10 h light/dark cycle. V1-leaf and V1-root samples were collected from maize seedlings that bore one leaf with the collar visible. V3-stalk samples were harvested from maize with three leaves having a visible collar. R1-leaf, R1-tassel, R1-silk and R1-ear samples were collected from maize that had begun flowering, i.e. when a silk was just visible outside the husks. Seeds were harvested from plants 50 to 60 days after pollination and dried before use. Germinating seeds were collected two days after germination. All tissues were harvested and immediately frozen in liquid nitrogen. At least two biological replicates were performed for each tissue.

### PARE library construction and data processing

RNA samples were extracted from maize tissues using TRIzol reagent (Molecular Research Center) according to the manufacturer’s instructions. The PARE libraries were constructed based on a previously described method (Zhai et al., 2014). PARE sequencing was performed on the Illumina HiSeq 2500 platform (Berry Genomics) to produce 50-bp single-end reads. After removing low-quality reads from the raw sequences, the adaptor sequences (TGGAATTCTCGGG) were prescinded using an in-house perl script. The CleaveLand4 pipeline (Addo-Quaye et al., 2009) was used to find sliced miRNA targets using maize B73 RefGen_v4 transcripts (Jiao et al., 2017) and all maize miRNA sequences from miRbase 22.1 (Kozomara et al., 2019) as input.

### RNA-seq library construction and data processing

mRNA libraries were constructed using NEBNext® UltraTM RNA Library Prep Kit for Illumina (NEB, USA) according to the instructions, and sequenced (150-bp paired-end reads) on the Illumina HiSeq 2500 platform at Berry Genomics (Beijing, China). Raw reads were processing by FastQC for quality control and mapped to the B73 RefGen_v4 genome (Jiao et al., 2017) using Hisat2 (Kim et al., 2015) after trimming the adaptor sequences.

### Small RNA-seq library construction and data processing

Small RNAs (15-to 40-nt) were isolated as described (He et al., 2019). The small RNA libraries were constructed using the NEBNext^®^ Multiplex Small RNA Library Prep Set for Illumina^®^ (NEB, E7300S) kit and sequenced on the Illumina HiSeq 2500 platform at Berry Genomics (Beijing, China) to generate 50-bp single-end reads. After removing the 3’-adaptor sequences (AGATCGGAAGAGC) using cutadapt (Martin, 2011), reads mapping to rRNA, tRNA, snoRNA and snRNA sequences were filtered using Bowtie2 (Langmead and Salzberg, 2012). The remaining reads were mapped to the B73 RefGen_v4 genome (Jiao et al., 2017) using ShortStack (Johnson et al., 2016), and the reads were counted and assigned to each miRNA to quantify the expression levels.

### Small RNA Northern blot

Total RNA (10 μg) was separated in a 15% polyacrylamide/urea gel after denaturation at 70°C for 10 min. The RNA was transferred onto a neutral nylon membrane (Hybond-NX, GE Healthcare) and crosslinked using 1-ethyl-3-(3-dimethylaminopropyl) carbodiimide hydrochloride (Sigma). Biotin-labelled probes were hybridized with sRNAs on the nylon membrane and stabilized streptavidin-horseradish peroxidase was used to detect the biotin signal.

### 5’-RACE

A Dynabeads mRNA purification kit (610-06) (Invitrogen) was used to isolate poly(A) RNA from total RNA (∼100 μg). Total RNA was extracted from different tissues of the inbred line B73 using TRIzol reagent (Molecular Research Center). After the 5’-RNA adaptor (5’-GTTCAGAGTGCTACAGTCCGACgtcagagctccga-3’) was ligated to the isolated poly(A) RNA, first-strand cDNA was synthesized using an RT primer (5’-CGAGCACAGAATTAATACGACTTTTTTTTTTTTTTTTTT-3’). An initial PCR was carried out using the 5’-RACE outer primer and gene-specific outer primer for cDNA synthesis. Nested PCR was then performed using the 5’-RACE inner primer, the gene-specific inner primer and 1/50 of the initial PCR reaction as template. The PCR products were sub-cloned into pCE2 TA (Vazyme, C601-02) and sequenced for further analysis.

### Dual luciferase reporter assays

To generate Pro35s::target site-LUC and Pro35s::mutated target site-LUC reporters for the dual-luciferase assays, the ∼60-bp fragments containing the target sites of putative target genes were inserted into the *BamH*I and *Nco*I sites of pGreenII-0800. Renilla LUC, under the control of CaMV 35S promoter, was used as the internal control (Hellens et al., 2005). The Pro35s::pre-miRNA effectors were created by cloning their precursor sequences into the *BamH*I and *Hind*III sites of pGreenII 62-SK. Transient dual-luciferase assays were conducted in 5-week-old *Nicotiana benthamiana* leaves using an *Agrobacterium tumefaciens*-mediated method. After growing for 48 h, LUC activities were measured using the TransDetect Double-Luciferase Reporter Assay kit (FR201-01) according to the manufacturer’s instructions. The relative ratio of firefly LUC to renilla LUC activity was calculated to represent the expression level of reporter genes.

### Primers and probes

All primers used for 5’-RACE and reporter constructs, as well as probes used for northern blots, are listed in Supplemental Dataset 6.

### Accession Numbers

The PARE-seq Illumina reads for all samples except R1-silk have been deposited in the Sequence Read Archive at the National Center for Biotechnology Information (http://www.ncbi.nlm.nih.gov/sra) under accession number PRJNA841081. RNA-seq and sRNA-seq reads for V3-stalk, R1-leaf, R1-tassel, R1-ear and R1-silk, as well as the PARE-seq for R1-silk, were deposited under accession number PRJNA520822.

## AUTHOR CONTRIBUTIONS

L. L., G. L. and J. H. conceived the project. G. L., J. H., C. X., and C. Y. performed the bioinformatics analysis. L. L., J. H. and C. X. designed and performed the experiments. L. L., C. X., and J. H. wrote the manuscript. B. M. and X. C. revised the manuscript.

## ACKNOWLEDGMENTS

We thank the Instrumental Analysis Center of Shenzhen University for technical assistance.

